# Eicosanoid signaling as a therapeutic target in middle-aged mice with severe COVID-19

**DOI:** 10.1101/2021.04.20.440676

**Authors:** Lok-Yin Roy Wong, Jian Zheng, Kevin Wilhelmsen, Kun Li, Miguel E. Ortiz, Nicholas J. Schnicker, Alejandro A. Pezzulo, Peter J. Szachowicz, Klaus Klumpp, Fred Aswad, Justin Rebo, Shuh Narumiya, Makoto Murakami, David K. Meyerholz, Kristen Fortney, Paul B. McCray, Stanley Perlman

## Abstract

Coronavirus disease 2019 (COVID-19) is especially severe in aged populations^1^. Resolution of the COVID-19 pandemic has been advanced by the recent development of SARS-CoV-2 vaccines, but vaccine efficacy is partly compromised by the recent emergence of SARS-CoV-2 variants with enhanced transmissibility^2^. The emergence of these variants emphasizes the need for further development of anti-SARS-CoV-2 therapies, especially in aged populations. Here, we describe the isolation of a new set of highly virulent mouse-adapted viruses and use them to test a novel therapeutic drug useful in infections of aged animals. Initially, we show that many of the mutations observed in SARS-CoV-2 during mouse adaptation (at positions 417, 484, 501 of the spike protein) also arise in humans in variants of concern (VOC)^2^. Their appearance during mouse adaptation indicates that immune pressure is not required for their selection. Similar to the human infection, aged mice infected with mouse-adapted SARS-CoV-2 develop more severe disease than young mice. In murine SARS, in which severity is also age-dependent, we showed that elevated levels of an eicosanoid, prostaglandin D2 (PGD_2_) and of a phospholipase, PLA_2_G2D, contributed to poor outcomes in aged mice^3,4^. Using our virulent mouse-adapted SARS-CoV-2, we show that infection of middle-aged mice lacking expression of DP1, a PGD_2_ receptor, or PLA_2_G2D are protected from severe disease. Further, treatment with a DP1 antagonist, asapiprant, protected aged mice from a lethal infection. DP1 antagonism is one of the first interventions in SARS-CoV-2-infected animals that specifically protects aged animals, and demonstrates that the PLA_2_G2D-PGD_2_/DP1 pathway is a useful target for therapeutic interventions. (Words: 254)

## TEXT

### Isolation of mouse-adapted virus

Mice are naturally resistant to infection with SARS-CoV-2, resulting from incompatibility between the viral surface (S) glycoprotein and mouse angiotensin converting enzyme 2 (mACE2)^5^. To address this incompatibility, human ACE2 (hACE2) has been provided by genetic manipulation of the mouse genome or exogenously using viral vectors. Transgenic expression of hACE2^6-9^, complete or partial replacement of mACE2 with hACE2 (hACE2-KI)^10^ or administration of viral vectors expressing hACE2 all sensitize mice for infection^11,12^. Alternatively, the spike protein (S) of SARS-CoV-2 has been altered using reverse genetics to enable binding to mACE2^13,14^. SARS-CoV-2 has been adapted to mice by targeting amino acids at positions 498/499 (Q498Y/P499T) or 501 (N501Y). The resulting viruses can infect mice, although the resulting infection is very mild. However, additional virus passage through mouse lungs results in increased virulence^15^. Mice infected with virulent mouse-adapted SARS-CoV or MERS-CoV (Middle East respiratory syndrome-CoV) have been similarly developed and are very useful for studies of pathogenesis, including in aged animals.

SARS, MERS, and COVID-19 show age-dependent increases in severity^1^. For example, no patient under the age of 24 years succumbed to SARS-CoV during the 2002-2003 epidemic, while 50% of those over 65 years died from the infection^16^. Parallel results were found in C57BL/6 (C57BL/6) mice infected with SARS. Young C57BL/6 mice were resistant to infection with mouse-adapted SARS-CoV while mice became progressively more susceptible beginning at about 5 months of age^4^. Poor survival correlated with a suboptimal T cell response, which resulted from delayed migration of respiratory dendritic cells (rDCs) to draining lymph nodes. We identified a single prostaglandin (PG), PGD_2_, that increased in the lungs during aging and was responsible for impaired DC migration^4^. We also showed that a single phospholipase A2, PLA_2_ group 2D (PLA_2_G2D), increased in mouse respiratory DCs during aging and was responsible for increased levels of PGD_2_, as well as other PGs in the aged mouse lung^3^. Given the similarities between SARS-CoV, MERS-CoV and SARS-CoV-2 in age-dependent increased disease severity, we postulated that PGD_2_ and PLA_2_G2D would have similar roles in aged mice infected with a virulent strain of mouse-adapted SARS-CoV-2.

To generate a virulent mouse-adapted SARS-CoV-2, we inserted the N501Y mutation into the SARS-CoV-2 genome (rSARS2-N501Y_P0_) using reverse genetics as previously described^17^. As expected, this mutation rendered mice susceptible to SARS-CoV-2 infection but BALB/c mice lost only a minimal amount of weight even after administration of 10^5^ PFU of virus. After 30 passages through mouse lungs, the virus became highly virulent such that 5000 PFU caused a lethal disease in young BALB/c mice (**Figure 1a**). We sequenced viruses after 10, 20 and 30 passages. Virus from these passages became progressively more lethal (**Figure 1a**). Focusing on changes in the S protein, passage 10 virus contained a Q498R mutation, which likely enhanced binding to mACE2^13^ (**Figure 2a**,**b**). By passage 20, a mutation at position 493 (Q493R) was detected. By passage 30, mutations in residues 417 and 484 (K417M, E484K) also arose. E484K does not appear to be required for mouse virulence since it was variably expressed by different isolates that had equivalent lethality (**Figure 2d**).

**Figure 1:**
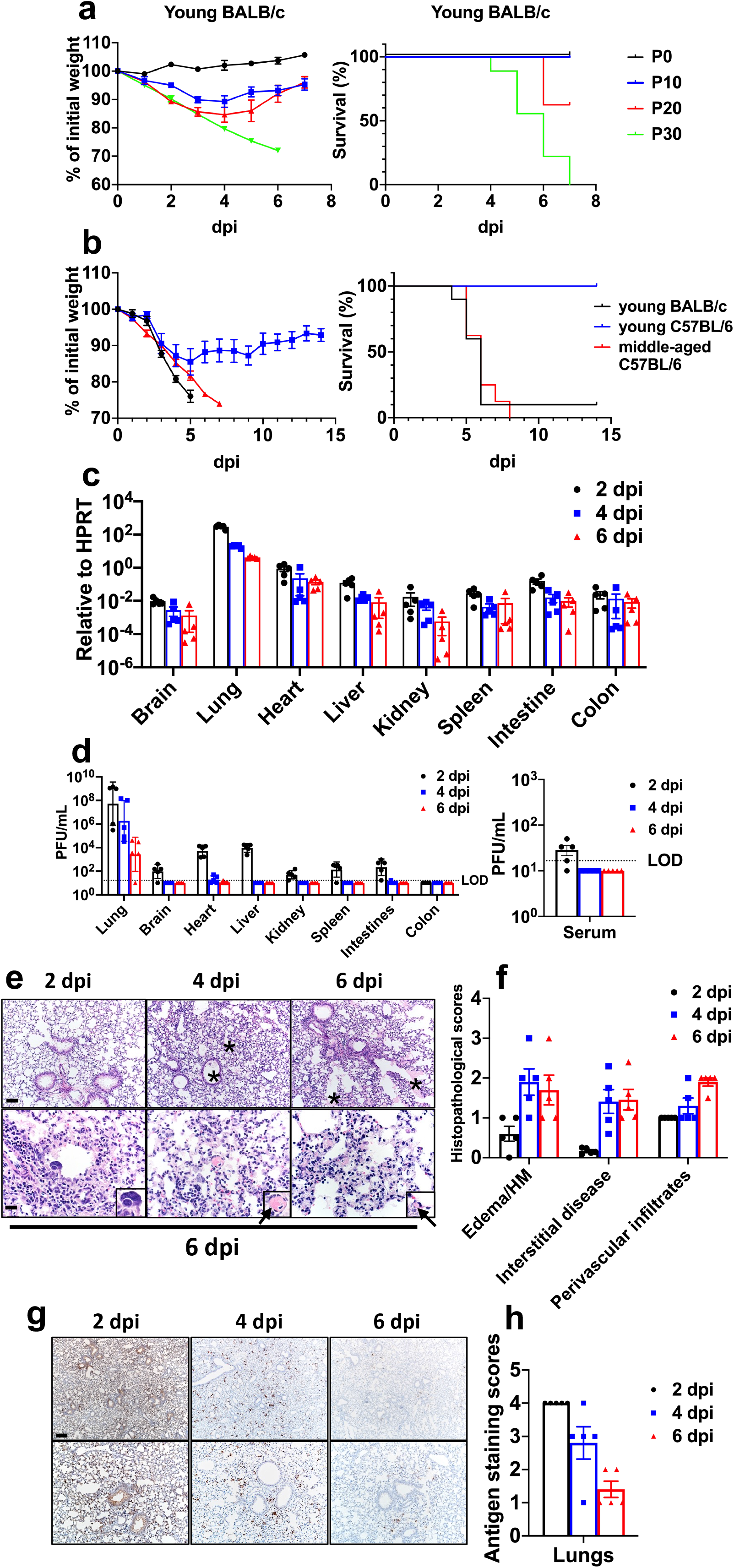
Clinical, virological and pathological disease in mice infected with SARS2-N501Y_MA30_. **a**, Percentage of initial weight and survival of young BALB/c mice infected with P0 (n=3, 10^5^ PFU), P10 (n=8, 9000 PFU), P20 (n=8, 9000 PFU) or P30 (n=9, 5000 PFU) of SARS-CoV-2 per mouse. “P” refers to the number of times that rSARS2-N501Y_P0_ was passaged in mouse lungs. **b**, Percentage of initial weight and survival of young BALB/c (n=10), young C57BL/6 (n=6) or middle-aged C57BL/6 (n=8) mice infected with 5000 PFU of SARS2-N501Y_MA30_ per mouse. Data in **a** and **b** are mean ± sem. **c, d**, Quantification of viral genomic RNA (**c**) and infectious viral titres detected by plaque assay (**d**) in different organs at 2, 4 and 6 days after infection with 5000 PFU of SARS2-N501Y_MA30_. (n=5 at all time points). Serum titers on 2, 4 and 6 dpi (n=5 at all time points) are shown in the right-hand panel. LOD, limit of detection. Data in **c** and **d** are geometric mean ± geometric SD **e-h:** Lungs from mice infected with 5000 PFU of SARS2-N501Y_MA30_ at 2, 4 and 6 dpi (n=5) were stained with H&E (**e**) or immunostained to detect SARS-CoV-2 nucleocapsid protein (**g**). **e**, Infected lungs exhibited airway edema (asterisks, top panels), multinucleated syncytial cells (inset in lower left panel), vascular thrombosis (arrows, insets in lower middle and right insets). Scale bar, 90 and 22 μm (top and bottom, respectively), H&E stain. **f**, Summary scores of lung lesions, as described in Methods (n=5). HM: hyaline membranes **g**, Lungs from mice infected with 5000 PFU of SARS2-N501Y_MA30_ at 2, 4 and 6 dpi (n=5) were immunostained to detect SARS-CoV-2 nucleocapsid protein. Scale bar 230 and 90 μm (top and bottom, respectively). **h**, Summary scores of nucleocapsid immunostaining, as described in Methods (n=5). Data in **f** and **h** are mean ± sem.

**Figure 2:**
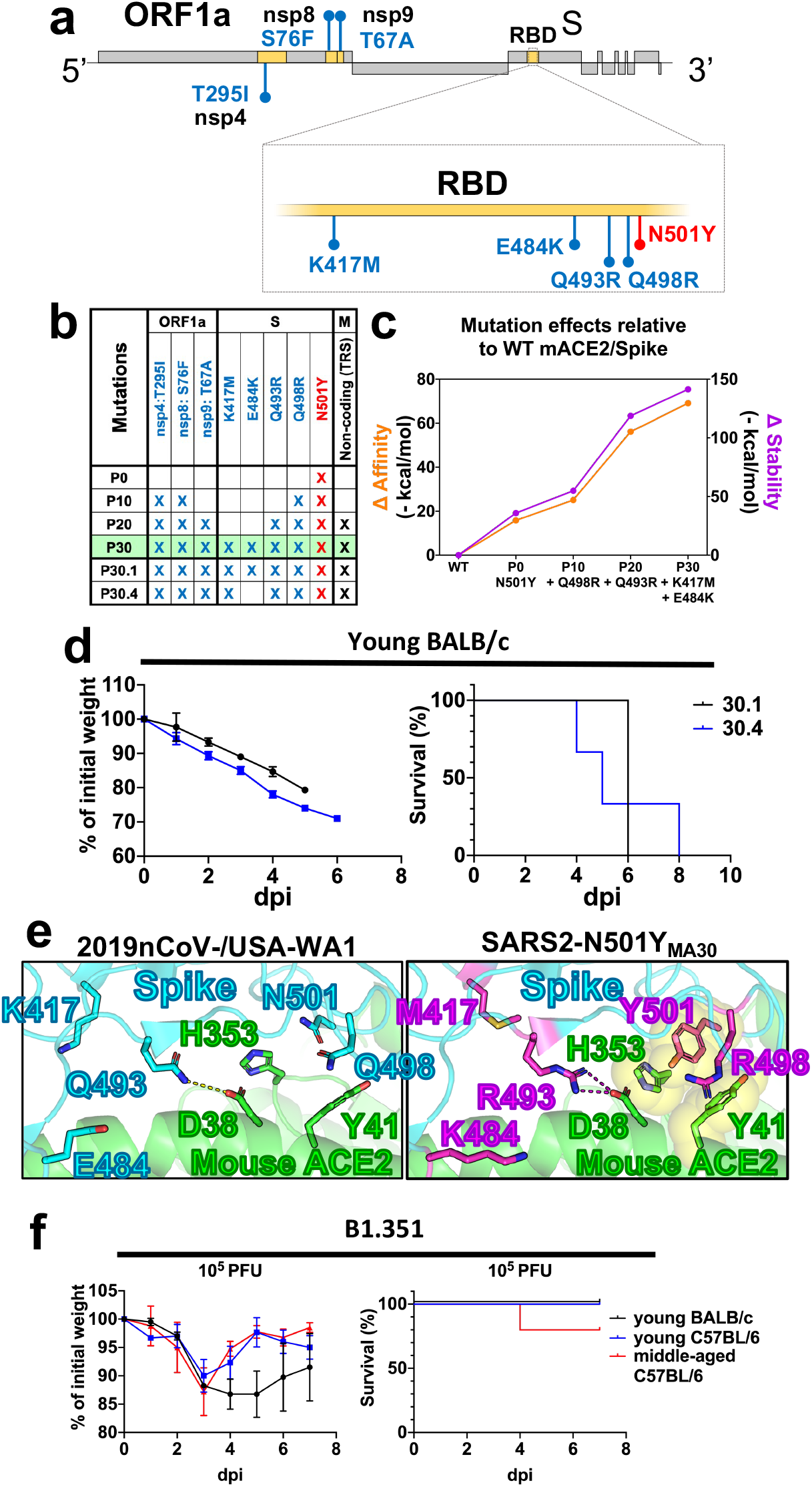
Analysis of mutations in SARS2-N501Y_MA30_ that arise during mouse adaptation. **a**, Schematic showing the genome of SARS-CoV-2. Initial N501Y mutation engineered into SARS-CoV-2 is shown in red. Viral proteins encoding mutations in SARS2-N501Y_MA30_ are shaded in yellow and the corresponding mutations are highlighted in blue. **b**, Summary of mutations emerged in different virus passages. P30 virus is denoted in green **c**, *In silico* modelling of the effects of mutations emerged in serial mouse passaging on the affinity and stability of the S protein complexes with mACE2. TRS-Transcriptional Regulatory Sequence. **d**, Percentage of initial weight and survival of young BALB/c mice infected with viruses from two distinct plaque isolates purified from P30 virus. Virus from plaque 30.1 (black) encodes an extra mutation (E484K) in the S protein compared to virus from plaque 30.4 (blue) (n=3 for both groups). **e**, Modeling of the receptor-binding interface between the S protein of 2019n-CoV/USA-WA1/2019 (left panel) and SARS2-N501Y_MA30_ (right panel) viruses and mouse ACE2 reveals critical interactions mediated by mutations emerged through serial passaging. **f**, Percentage of initial weight and survival of young BALB/c, and young or old C57BL/6 mice infected with 10^5^ PFU of B1.351. (n=4 for young BALB/c; n=3 for young C57BL/6; n=5 for middle-aged C57BL/6). Data in **b** and **f** are mean ± sem.

*In silico* modeling of the interaction between mouse ACE2 and the spike glycoprotein showed that Y501 in the spike allowed for pi-pi interactions and hydrophobic interactions to occur with Y41 and H353 in mACE2, increasing the affinity and stability of the complex (**Figure 2c**,**e**). Two additional spike mutations (Q498R and Q493R) further increased affinity and stability, mainly driven by R493 which can create a salt bridge with D38 in mACE2 (previously a hydrogen bond between Q493 in spike and D38 in mACE2). Mutations to other positively charged amino acids in residue 493 of the spike have also been described in another mouse-adapted SARS-CoV-2^15^ suggesting the importance of this salt bridge to D38 in mACE2. The contribution of R498 in the spike is likely due to an increase in the buried solvent-accessible surface area of this residue (from 80% to 99%), which improves the overall stability of the molecule and in turn could increase the affinity for ACE2. Residues K417 and E484 in the spike do not interact directly with ACE2, and have been reported to have minimal effects on hACE2 binding^18^. E484K and various mutations in residue 417 have been associated with VOC B.1.351 (E484K + K417N) and P.1 (E484K + K417T). E484K and K417N have been shown to decrease antibody-mediated neutralization^19^ and therefore have been hypothesized to emerge in humans as an evolutionary advantage to evade immune responses. Here we show that E484K and mutations in K417 also occur in the absence of an adaptive immune response, offering an alternative explanation for the evolutionary pressures that drive the appearance of mutations in residues 417 and 484. Previously published models of non-lethal murine SARS-CoV-2 have also reported mutations identical to the ones described here after 30 mouse lung passages^15,20,21^ demonstrating the advantage of this particular combination of mutations in driving murine lethality.

Mutations in three nonstructural proteins, nsp4, nsp8, and nsp9 were detected at passage 20. Regarding these, T295I in nsp4 and T67A in nsp9 have been described in other models of SARS-CoV-2 mouse adaptation that caused moderate disease but were not lethal in young BALB/c mice^15^, suggesting these mutations might be important in driving virulence but are insufficient to cause death. T67A in nsp9 has also been described in a lethal mouse model of SARS-CoV^22^, suggesting that this mutation might be important for mouse adaptation not only in SARS-CoV-2 but in other coronaviruses as well. To assess the importance of Q493R and the nsp mutations in virulence, we infected young BALB/c and middle-aged C57BL/6 mice with the B.1.351 strain, which contains K417N, E484K, and N501Y. Infection with a high dose of virus (10^5^ PFU) resulted in modest weight loss and 20% lethality, indicating that one or more of the mutations detected in nsp4, nsp 8, nsp9 and Q493R were required for mouse virulence (**Figure 2f**).

Virus was then plaque purified three times and individual plaques were sequenced. One plaque was chosen for further study and propagated (SARS2-N501Y_MA30_). Of note, SARS2-N501Y_MA30_ grew less well than rSARS2-N501Y_P0_ in human airway epithelial (Calu-3) cells, showing that increased virulence in mice did not result in increased replication in human cells (**Extended Data Figure 1a**).

### Clinical and pathological characterization of mouse adapted viruses

To further assess the virulence of SARS2-N501Y_MA30_, we infected 6-10 week old BALB/c and C57BL/6, and 6-9 month old C57BL/6 mice intranasally with 5000 PFU (**Figure 1b**). 5000 PFU SARS2-N501Y_MA30_ caused a lethal infection in young BALB/c mice and middle-aged C57BL/6 mice but only modest weight loss in young C57BL/6 mice. For most of our subsequent experiments, we infected young BALB/c or middle-aged C57BL/6 mice with SARS2-N501Y_MA30_. After infection of young BALB/c mice with 5000 PFU, viral genomic RNA was detected in most organs at all time points, although levels were much higher in the lungs (**Figure 1c**). Infectious virus was detected in most organs at day 2 but by 4 and 6 days post infection (dpi), it was detected only in the lungs, and to a lesser extent, the heart (**Figure 1d**). Consistent with the presence of virus in organs, we also detected virus in the blood of 4/5 mice at 2 dpi (**Figure 1d, right**). To extend these results, we analyzed histological samples for evidence of pathological changes and virus antigen (**Figure 1e-h**). We detected pathological changes in the lungs and nasal cavity at all times, but not in other organs (**Figure 1e**,**f, Extended Data Figure 2**). Pathological findings in the lung included perivascular, peribronchial and interstitial infiltration and alveolar edema. We also detected occasional pulmonary vascular thrombi, which are prominent findings in patients with severe COVID-19^23^ and rare multinuclear syncytia (**Figure 1e**, insets, lower panels). In the sinonasal cavity, we detected evidence of infection of the olfactory and respiratory epithelium, with prominent infection of sustentacular cells (**Extended Data Figure 1b**,**c**). Unlike SARS-CoV-2 infection of K18-hACE2 mice^9^, there was no pathological or immunohistochemical evidence of infection in the brain (**Extended Data Figure 1d**,**e)**.

### Immunological response to SARS2-N501Y_MA30_ infection

To further characterize the immunological response in the lungs to SARS2-N501Y_MA30_ infection, we measured levels of several cytokines, chemokines and inflammation-associated molecules in BALB/c lungs at days 2, 4, and 6 p.i. Mice were infected with 1000 PFU in these assays to ensure that most survived until 6 dpi. Increases in IFNs (IFNα, IFNβ, IFNγ, IFNλ), ISGs (ISG15, OAS1), cytokines (IL-1α, IL-1β, IL-6, IL-8, IL-12, IL-15), chemokines (CCL2, CCL5, CXCL2, CXCL9, CXCL10) and CCR7, a chemokine receptor required for cell migration to draining lymph nodes, were observed over the course of the infection, with peak levels of IFNα, IFNβ, and IFNλ prominent at 2 dpi. **(Extended Data Figure 3a)**. Notably, levels of most of these pro-inflammatory molecules remained elevated throughout the infection, as also occurs in the blood of COVID-19 patients^24^. To determine the character of the cells infiltrating the lungs observed on pathological examination, we immunophenotyped the infiltrating cells at 4 and 6 dpi. **(Extended Data Figure 3b**,**e)**. Over time, there was a gradual increase in the numbers of infiltrating CD11b^+^ cells (macrophages, monocytes, and neutrophils) **(Extended Data Figure 3b)**. Virus-specific CD4 and CD8 T cells were also detected in the lungs at day 6 p.i. These cells expressed IFNγ and TNF in response to stimulation with pools of peptides covering the SARS-CoV-2 structural proteins (spike, membrane, and nucleocapsid) **(Extended Data Figure 3c**,**d)**. The S, N, and M proteins all elicited CD4 and CD8 T cell responses, with the S protein responsible for the largest responses in both cell types. Lymphopenia is a hallmark of severe COVID-19^25^ and was detected in SARS2-N501Y_MA30_-infected middle-aged mice **(Extended Data Figure 4)**.

**Figure 3:**
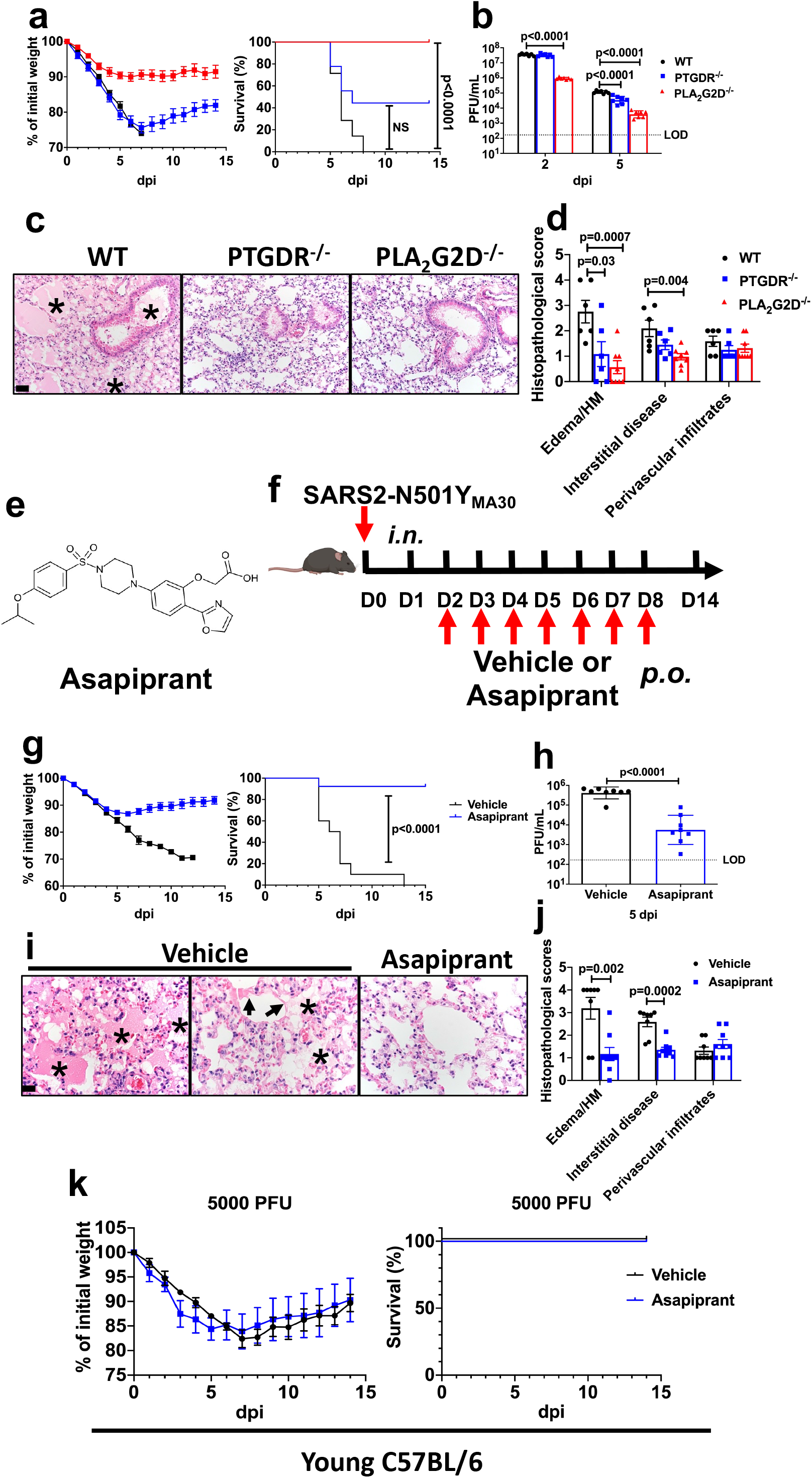
Absence of PLA_2_G2D or DP1 expression, or blocking PGD_2_/DP1 signaling inhibits SARS2-N501Y_MA30_ infection in vivo. **a**, Percentage of initial weight and survival of middle-aged WT (n=8), PTGDR^-/-^ (n=9) or PLA_2_G2D^-/-^ (n=9) C57BL/6 mice infected with 5000 PFU of SARS2-N501Y_MA30_ per mouse. **b**, Infectious viral titres detected by plaque assay in the lungs of middle-aged WT, PTGDR^-/-^ or PLA_2_G2D^-/-^ mice at 2 (n=7), and 5 dpi (n=7) with 5000 PFU of SARS2-N501Y_MA30_ per mouse. LOD, limit of detection. *P* values determined by log-rank (Mental-Cox) (**a**) and two-tailed Student’s t test (**b**). \**P*<0.0001, PLA_2_G2D^**-/-**^ vs WT. NS, not significant (**a**). \**P*<0.0001, PLA_2_G_2_D^**-/-**^ vs WT, 2dpi; *P<0.0001, PTGDR^-/-^ vs WT and PLA_2_G2D^-/-^ vs WT, 5 dpi (**b**). **c**, Infected lungs at 5 dpi from WT mice exhibited common features of diffuse alveolar damage including edema (*), and cellular infiltration/interstitial thickening. Infected lungs at 5 dpi from PTGDR^-/-^ or PLA_2_G2D^-/-^ mice exhibited significant reduction in edema and interstitial disease, H&E stain. Scale bar 40 μm. **d**, Summary scores of lung lesion, as described in Methods (n=6 for WT; n=6 for PTGDR^-/-^; n=8 for PLA_2_G_2_D^**-/-**^). *P* values determined by two-tailed Student’s t test. \**P*=0.03, PTGDR^-/-^ vs WT and \**P*=0.0007, PLA_2_G2D^-/-^ vs WT, Edema/HM; \**P*=0.004, PLA_2_G2D^-/-^ vs WT, Interstitial disease. **e**, Structure of asapiprant. **f**, Schematic showing experimental design for results shown in panels **g-k** (image of the mouse shown was created with BioRender.com). Vehicle or asapiprant were administered to infected mice orally from 2-8 dpi to middle aged (**g-j**) or young (**k**) C57BL/6 mice infected with 5000 PFU of SARS2-N501Y_MA30_. **g**, Percentage of initial weight and survival of vehicle-(n=10) or asapiprant-treated (n=12) middle-aged mice infected with 5000 PFU of SARS2-N501Y_MA30_ per mouse. *P* values determined by two-tailed Student’s t test. \**P*<0.0001, asapiprant-vs vehicle-treated group. **h**, Infectious viral titres detected by plaque assay in the lungs of vehicle-(n=8) or asapiprant-treated (n=8) middle-aged mice at 5 dpi infected with 5000 PFU of SARS2-N501Y_MA30_ per mouse. *P* values determined by two-tailed Student’s t test. \**P*<0.0001, asapiprant-vs vehicle-treated group. **i**, Vehicle-treated group exhibited widespread edema (asterisks) with occasional hyaline membranes (arrows) whereas these features were uncommon in asapiprant-treated mice at 5 dpi, H&E stain. Scale bar 20 μm. **j**, Summary scores of lung lesions, as described in Methods (n=8 for vehicle-treated mice; n=9 for asapipant-treated mice). *P* values determined by two-tailed Student’s t test. \**P*=0.002, asapiprant-vs vehicle-treated group, Edema/HM; \**P*=0.0002, asapiprant-vs vehicle-treated group, Interstitial disease. HM: hyaline membranes. **k**, Percentage of initial weight and survival of vehicle- (n=4) or asapiprant-treated (n=4) young mice. Data in **a, d, g, j, k** are mean ± s.e.m. Data in **b, h** are geometric mean ± geometric SD.

### Role of PGD_2_/DP1 signaling and PLA_2_G2D expression in disease severity infected mice

Since we previously identified roles for PGD_2_/DP1 signaling and PLA_2_G2D expression by respiratory DCs in enhanced disease in SARS-CoV-infected middle-aged mice^3,4^, we next infected 8-12 month old *PLA*_*2*_*G2D*^*-/-*^ and *PTGDR*^*-/-*^ (lacking DP1 expression) mice with 5000 PFU, a lethal dose of SARS2-N501Y_MA30_. Lethality was completely or partially abrogated in the absence of PLA_2_G2D or DP1 expression, respectively **(Figure 3a)**. In both cases, the kinetics of virus clearance was enhanced, likely contributing to diminished disease **(Figure 3b)**. Further, many of the pathological changes observed in the lungs of infected middle-aged C57BL/6 mice, such as extensive inflammatory infiltration and edema, were not present in either *PLA*_*2*_*G2D*^*-/-*^ and *PTGDR*^*-/-*^ mice **(Figure 3c**,**d)**. Together, these results show that PLA_2_G2D, associated with anti-inflammatory effects in mice infected with SARS-CoV, also contributes to poor outcomes in SARS-CoV-2 infected mice. The results also demonstrate that these protective effects were at least partially mediated through the PGD_2_/DP1 axis.

### Antagonizing PGD_2_/DP1 signaling with asapiprant

While several drugs with inhibitory activity against PLA_2_ molecules have been described^26^, none are specific for murine PLA_2_G2D. However, asapiprant (BGE-175) ([2-(Oxazol-2-yl)-5-(4-{4-[(propan-2-yl)oxy]phenylsulfonyl} piperazine-1-yl)phenoxy] acetic acid), a potent and specific antagonist of human PGD_2_/DP1 signaling, has been previously described (**Figure 3e**)^27^. Asapiprant is a potent and selective antagonist of PGD_2_-signaling via the DP1 receptor with a mean K_i_ of 0.44 nM. Asapiprant affinity for DP1 was 300 to >15000-fold higher than for any other prostanoid receptor tested^27^. We administered asapiprant to mice by once daily oral gavage (30 mg/kg), starting two days after infection with 5000 PFU of SARS2-N501Y_MA30_ and for 6 days thereafter **(Figure 3f)**. For these experiments, we used middle-aged C57BL/6 mice. We chose to start treatment at two days after infection because it is relevant for possible use in symptomatic patients. In middle-aged mice infected with a lethal dose of SARS2-N501Y_MA30_, asapirant reduced mortality from 100% to <10%, with concomitant effects on weight loss **(Figure 3g)**. As expected based on the studies of *PLA*_*2*_*G2D*^*-/-*^ and *PTGDR*^*-/-*^ mice described above, drug treatment resulted in faster kinetics of virus clearance **(Figure 3h)** and diminished pathological changes in the lungs **(Figure 3i**,**j)**. In contrast, treatment with the drug did not reduce the weight loss observed in young SARS2-N501Y_MA30_-infected C57BL/6 mice **(Figure 3k)**.

## Discussion

We describe a mouse-adapted SARS-CoV-2, isolated from mouse lungs after 30 serial passages. Unlike previously described mouse-adapted viruses, this strain is highly virulent in young and old mice, and infects only the lungs and not the brain. Further, it contains a set of mutations that are also observed in the B.1.351 and P.1 strains (K417N/T, E484K and N501Y) although K417N/T is replaced by K417M in SARS2-N501Y_MA30_. All three amino acids are located in the receptor binding domain and E484 and N501 are hotspots for recognition by neutralizing antibodies^19^. N501Y enhances binding to ACE2 and transmission in hamsters^28^, while K417N and E484K do not appear to enhance binding to ACE2^18^. Our results show that the selection of K417M (and by extension, presumably K417N) and E484K does not result solely from evasion of the antibody response since they arose in mice in the absence of any immune pressure. Infection of mice with B1.351 resulted in mild disease, indicating that changes outside of the S protein were critical for mouse virulence. T295I, located in the transmembrane domain of nsp4 and Q493K in the receptor binding domain of S were identified in another variant of mouse-adapted SARS-CoV-2^15^ so that mutations at these sites are prime candidates for conferring virulence in mice.

Our results identify a key role for PGD_2_/DP1 signaling in enhanced COVID-19 disease severity during aging and show that delivery of a DP1 inhibitor at 2 dpi, a time comparable to when drug might be used in patients, converted a lethal to a sublethal infection. Asapiprant has been used in previous studies of allergic rhinitis and asthma. In a rat model of asthma, oral asapiprant reduced inflammatory cell numbers in BAL fluid and reduced the increase in airway hyperresponsiveness (AHR)^27^. Asapiprant has been evaluated in 18 clinical studies (Phase 1, 2, 3) (2000 patients and 200 healthy volunteers) conducted in the US, UK, France, and Japan for the treatment of allergic rhinitis, alone or in combination with cetirizine, an anti-histamine^29^. Once-daily dosing of asapiprant for up to 4 weeks was considered safe and well tolerated. Decreased airway resistance after PGD_2_ challenge and decreased rhinitis symptom scores after allergen exposure were demonstrated in these studies, consistent with on-target pharmacodynamics^29^. Asapiprant is now being used in randomized, placebo-controlled clinical trials of hospitalized COVID-19 patients who are at risk for respiratory failure (https://clinicaltrials.gov/ct2/show/NCT04705597). Together, our results show that infection of mice with SARS2-N501Y_MA30_ recapitulates many of the findings observed in patients with severe COVID-19 and also identify PGD_2_/DP1 signaling and PLA_2_G2D as useful targets for therapy in aged individuals.

## Online content

Any methods, Nature Research reporting summaries, source data, extended data, supplementary information, acknowledgments, peer review information; details of author contributions and competing interests; and statements of data and code availability are available at https://doi.org/.

## METHODS

### Generation of SARS-CoV-2 encoding N501Y mutation in the S Protein

p-BAC SARS-CoV-2 carrying the sequence of isolate Wuhan-Hu-1 (WT-SARS-CoV-2 BAC) was a gift from Drs. Sonja Zuniga and Luis Enjuanes (CNB-CSIC, Madrid, Spain). The only difference between the Wuhan-Hu-1 and 2019n-CoV/USA-WA1/2019 strains is one amino acid change in ORF8 and 2 silent mutations (in ORF1a and ORF1b). Two silent mutations were introduced into the BAC clone to enable differentiate from wild type SARS-CoV-2 (A20085G and G26840C). p-BAC SARS-CoV-2 with a N501Y mutation in the spike (N501Y-SARS-CoV-2 BAC) was generated using a lambda red recombination system with I-SceI homing endonuclease as previously described^17^. In brief, forward and reverse primers with overlapping SARS-CoV-2 spike sequence with the N501Y mutation followed by sequence complementary to the target plasmid (pEP-KanS) were synthesized (Invitrogen). A PCR fragment with overlapping ends carrying SARS-CoV-2 sequence flanking the Kanamycin resistance marker was amplified with the primers described using pEP-KanS as template. The PCR fragment was purified using gel purification kit (Invitrogen). Purified PCR fragment was electroporated into GS1783 strains of E. coli carrying p-BAC SARS-CoV-2. Successful recombinants were selected with kanamycin resistance. The kanamycin marker was removed by arabinose induction of I-SceI cleavage followed by homologous recombination of the overlapping ends. Successful recombinants were selected by replica plating for the loss of kanamycin resistance. Recombinants were purified and sequenced to confirm the introduction of N501Y mutation into the BAC clone. The primer sequences used were; Forward: 5’-ACCTTGTAATGGTGTTGAAGGTTTTAATTGTTACTTTCCTTTACAATCATAT GGTTTCCAACCCACT*<u>TAT</u>*GGTGTTGGTT**AGGATGACGACGATAAGTAG**-3’.

Reverse: 5’-CATGTAGAAGTTCAAAAGAAAGTACTACTACTCTGTATGGTTGGTAACCAA CA CCATTAGTGGGTTGGAAACCAT**CAACCAATTAACCAATTCTGATTAG**-3’. N501Y mutations are italicized and underlined; sequences complementary to pEP-KanS are bolded. 2 μg of N501Y-SARS-CoV-2 BAC were transfected into Vero E6 (ATCC) with Lipofectamine 3000 (Invitrogen) in a 6-well plate according to manufacturer’s protocol. Cells were monitored daily for cytopathic effects (CPE). Cultures were harvested when CPE was >50% by freezing at -80°C. rSARS2-N501Y_P0_ was further passaged in Calu-3 cells in DMEM supplemented with 20% FBS. Virus titer was determined by plaque assay.

### Mice, cells and virus

6-to-10-week- or 6-month-old BALB/c and C57BL/6 mice were obtained from Charles River Laboratories. 2019n-CoV/USA-WA1/2019 (Accession number: MN985325.1) and 20H/501Y.V2 (B.1.351, BEI catalogue number NR-54009) were obtained from BEI and passaged in Calu-3 2B4 cells^30^. Calu-3 2B4 cells were grown in Dulbecco’s modified Eagle’s medium (DMEM, GIBCO, Grand Island, NY) supplemented with 20% FBS. Vero E6 cells (ATCC CRL-1586) were grown in DMEM supplemented with 10% fetal bovine serum (FBS). All viruses were sequenced after propagation and found to match the input strain.

### Serial in vivo passaging of virus in mice

Serial blind passage of rSARS2-N501Y_P0_ through mouse lungs was performed in 8-to-10-week-old BALB/c mice. In brief, two mice were each inoculated with 50 μl of the virus intranasally at each passage. At 2/3 dpi, mice were sacrificed, and lungs were pooled, homogenized in PBS and used to infect naïve mice. After 30 passages, virus was obtained and plaque purified three times using Vero E6 cells. Several isolates were obtained and a single one was chosen for further study (SARS2-N501Y_MA30_). SARS2-N501Y_MA30_ was further propagated in Calu-3 2B4 cells.

### Mouse infection

Mice were anesthetized with ketamine-xylazine and infected intranasally with the indicated amount of virus in a total volume of 50 μl of DMEM. Animal weight and health were monitored daily. All experiments with SARS-CoV-2 were performed in a Biosafety Level 3 (BSL3) Laboratory at the University of Iowa. All animal studies were approved by the University of Iowa Animal Care and Use Committee and meet stipulations of the Guide for the Care and Use of Laboratory Animals.

### Virus titer by plaque assay

At the indicated times, mice were euthanized and transcardially perfused with PBS. Organs were harvested and homogenized prior to clarification by centrifugation and titering. Virus or tissue homogenate supernatants were serially diluted in DMEM. 12 well plates of VeroE6 cells were inoculated at 37°C in 5% CO_2_ for 1 h and gently rocked every 15 min. After removing the inocula, plates were overlaid with 0.6% agarose containing 2% FBS. After 3 days, overlays were removed, and plaques visualized by staining with 0.1% crystal violet. Viral titers were quantified as PFU/mL tissue.

### Collection of whole blood/serum

Mice were anesthetized by intraperitoneal injection of ketamine-xylazine. Blood was collected through retro-orbital bleed with a capillary tube (Fisher Scientific). Blood was allowed to clot at room temperature for 30 minutes. Serum was clarified by centrifugation and transferred to a new tube for storage at -80°C. For collection of whole blood, heparinized capillary tubes were used (Fisher Scientific).

### Histology and Immunohistochemistry

Mice were anesthetized by intraperitoneal injection of ketamine-xylazine and perfused transcardially with PBS. Tissues were fixed in zinc formalin. For routine histology, tissue sections (∼4 µm each) were stained with hematoxylin and eosin. Tissues were evaluated for the presence of edema or hyaline membranes (0-4) using distribution-based ordinal scoring: 0 – none; 1 - <25%; 2 - 26-50%; 3 - 51-75% and >75% of tissue fields. Perivascular lymphoid inflammation was evaluated by severity-based ordinal scoring: 0 – absent; 1-minor, solitary to loose infiltration of cells; 2 – moderate, small to medium aggregates; and 3 – severe, robust aggregates that are circumferential around vessels and can extend into adjacent parenchyma. Interstitial scores were evaluated using a modified H-Score for the presence of interstitial disease: 0 – none; 1 – minor, increased cellularity in septa; 2-moderate, cellular infiltrates with some thickened septa and uncommon inflammatory cells extending into lumen; and 3 – severe, cellular infiltrates in thickened septa and filling in air space lumens with atelectasis and / or evidence of diffuse alveolar damage (edema or hyaline membranes). For each score, this was multiplied by the % of lung affected, then summed for each lung and divided by 100 to yield a final score of 0-3. For SARS-CoV-2 antigen detection, slides were incubated with blocking reagent (10% normal goat serum x 30 minutes) followed by rabbit monoclonal antibody against SARS-CoV-2 N protein (1:20,000 dilution x 60 minutes, #40143-R019, Sino Biological US Inc., Wayne, PA, USA), then incubated with Rabbit Envision (Dako) and diaminobenzidine (Dako) as chromogen. Lung tissues sections were evaluated by a boarded veterinary pathologist familiar with the model using the post-examination method of masking^31^. IHC was ordinally scored on the % distribution of staining in the tissues: 0 – absent, 1 – 0-25%; 2 – 26-50%; 3 – 51-75%, and 4 - >75% of tissue.

### Lung cell preparation, antibodies for flow cytometric analysis

Animals were anesthetized with ketamine-xylazine and perfused transcardially with 10 mL PBS. Lungs were removed, minced, and digested in HBSS buffer consisting of 2% fetal calf serum, 25 mM HEPES, 1 mg/mL collagenase D (Roche) and 0.1 mg/mL DNase (Roche) at 37°C for 30 minutes. Single-cell suspensions were prepared by passage through a 70 μm cell strainer. Cells were enumerated with a Scepter 2.0 cell counter (MilliporeSigma). Whole blood was treated with ACK lysis buffer for 1 minute. Cells were washed and pelleted. Cells were then washed and blocked with 1 μg α-CD16/α-CD32 (2.4G2) antibody at 4 °C for 20 minutes and surface stained with the following antibodies at 4°C for 30 minutes: V450 α-mouse CD45 (clone 30-F11; Cat. No.: 560501) APC α-mouse CD220 (clone RA3-6B2, Cat. No.: 553092); APC/Cyanine 7 α-mouse CD3e (clone 145-2C11, Cat. No.: 100330); APC/Cyanine 7 α-mouse CD11c (clone HL3; Cat. No.: 561241); FITC α-mouse Ly6G (clone 1A8; Cat. No.: 127606); PE α-mouse CD11b (clone M1/70; Cat. No.: 101208); PE/Cyanine 7 α-mouse CD8 (clone 53-6.7; Cat. No.: 100722); PerCP/Cyanine 5.5 α-mouse CD4 (clone RM4.5; Cat. No.: 550954); FITC α-mouse Ly6G (1A8; Cat. No.: 127606); PerCP/Cyanine 5.5 α-mouse Ly6C (clone HK1.4; Cat. No.: 128012); CD 64 (clone X54-5/7.1; Cat. No.: 139304) Cells were washed, fixed and permeabilized with Cytofix/Cytoperm (BD Biosciences).

For intracellular cytokine staining (ICS), lymphocytes were cultured in 96-well dishes at 37°C for 5-6 h in the presence of 2 μM peptide pool and brefeldin A (BD Biosciences). Cells were then labeled for cell-surface markers, fixed/permeabilized with Cytofix/Cytoperm Solution (BD Biosciences), and labeled with α-IFN-γ and α-TNF antibody. All flow cytometry data were acquired using a BD FACSVerse and analyzed with FlowJo software.

### RNA isolation and RT-qPCR

Total RNA was extracted from tissues using TRIzol (Invitrogen) or a Direct-zol RNA Miniprep kit (Zymo Research) according to the manufacturer’s protocol. Following DNase treatment, 200 ng of total RNA was used as a template for first strand cDNA using a High-Capacity cDNA Reverse Transcription Kit (Applied Biosystems). The resulting cDNA was subjected to amplification of selected genes by real-time quantitative PCR using Power SYBR Green PCR Master Mix (Applied Biosystems). Average values from duplicates of each gene were used to calculate the relative abundance of transcripts normalized to HPRT and presented as 2^-ΔCT^. The primers used for cytokine and chemokines were previously reported^32^. For detection of viral genomes, the following primers were used to amplify the genomic RNA for the N protein: 2019-nCoV_N1-F: 5’-GAC CCC AAA ATC AGC GAA AT-3’; 2019-nCoV_N1-R: 5’-TCT GGT TAC TGC CAG TTG AAT CTG-3’. The following primers were used to amplify the subgenomic RNA for the E protein: F: 5’-CGATCTCTTGTAGATCTGTTCTC-3’; R: 5’-ATATTGCAGCAGTACGCACACA-3’.

### Viral whole genome sequencing

Samples were treated with TRIzol reagent for viral inactivation and RNA extraction. Sequencing libraries were prepared using the sequencing protocol (v2) and primer pools (v3) from the ARTIC network (dx.doi.org/10.17504/protocols.io.bdp7i5rn). Reads were sequenced on a MinION sequencer from Oxford Nanopore Technologies and consensus sequences were generated using Medaka following the ARTIC network nCoV-2019 bioinformatics protocol v1.1.0 (https://artic.network/ncov-2019/ncov2019-bioinformatics-sop.html). 249,944 average reads +/-82,509 (standard deviation) were obtained per sample. After quality filtering and alignment, average sequencing depth was >200X per amplicon for each sample. The consensus sequence of P10 and P20 virus from mouse lung homogenates contained areas with low sequencing depth; in order to accurately cover these gaps, we amplified the missing regions using individual PCR reactions and performed Sanger sequencing on the amplicons.

### In silico structural modelling and analysis

A homology model for mouse ACE2 (NP_001123985.1) was generated using YASARA and docked into the crystal structure of complexed human ACE2 and spike RBD (PDB: 6M0J). Mutation calculations were performed on the mACE2-Spike complex in Bioluminate (Schrodinger Release 2020-4) by first using the Protein Preparation Wizard and then the Residue Scanning module. Stability and affinity calculations were performed optimizing for the affinity and backbone minimization was used with a cutoff of 5 Å. Figures were generated in PyMOL. Software used was installed and configured by SBGrid^33^.

### Treatment with asapiprant

For asapiprant treatment, 7-8 month old, male and female C57BL/6 mice were infected with SARS2-N501Y_MA30_ (5000 PFU). Mice were treated daily from days 2-8 after infection with 0.2 ml vehicle (0.5% Carboxymethylcellulose Sodium (Na-CMC), Sigma), or 30 mg/kg asapiprant (BioAge) diluted in 0.2 ml vehicle. Drug was administered via oral gavage.

### Statistical analysis

Differences in mean values between groups were analysed by ANOVA and Student’s *t*-tests and differences in survival were analysed by log-rank (Mantel–Cox) tests using GraphPad Prism 8. All results are expressed as mean ± standard error of the mean and were corrected for multiple comparisons. *P* < 0.05 was considered statistically significant. *\*P* ≤ 0.05, *\*\*P* ≤ 0.005, *\*\*\*P* ≤ 0.0005, *\*\*\*\*P* ≤ 0.0001.

## AUTHOR CONTRIBUTIONS

L-Y. R. W. isolated mouse adapted virus, designed the study and experiments, and contributed to data interpretation and manuscript preparation. J.Z., K.L. contributed to experimental design, data interpretation, and manuscript preparation. K.W. contributed to experimental design, manuscript writing and data interpretation. M.E.O. performed the sequence analyses, modeling and manuscript preparation. N.J.S. performed *in silico* modeling and analysis. A.A.P. performed the sequence analyses. P.S. performed sequence analysis. K.K., F.A., J.R. contributed to experimental design, manuscript preparation and data interpretation. S.N., M.M provided critical reagents. D.K.M. analyzed and scored the pathological sections. K.F. contributed to manuscript preparation. P.B.M. designed experiments and contributed to data interpretation and manuscript preparation. S.P. designed and coordinated the study, designed experiments and contributed to data interpretation, data presentation and manuscript preparation.

## DATA AVAILABILITY

The data supporting the findings of this study are documented within the paper and are available from the corresponding authors upon request. Correspondence and requests for materials should be addressed to: Stanley Perlman or Paul B. McCray, Jr.

## ACKNOWLEDGMENTS

This work is supported in part by grants from the National Institutes of Health USA (NIH) (P01 AI060699 (SP, PBM) and RO1 AI129269 (SP) and BIOAGE Labs (SP). The Pathology Core, is partially supported by the Center for Gene Therapy for Cystic Fibrosis (NIH Grant P30 DK-54759), and the Cystic Fibrosis Foundation. PBM is supported by the Roy J. Carver Charitable Trust. We thank Dr. Michael Gelb (University of Washington) for PLA_2_G2D^-/-^ mice.

## EXTENDED DATA FIGURE LEGENDS

**Extended Data Figure 1. Infection of human respiratory cells and distribution of SARS2-N501Y**_**MA30**_ **antigen in sinonasal cavity and brain**.

**a**, Quantification of genomic RNA (gRNA), subgenomic RNA (sgRNA) and virus titers in Calu-3 cells at the indicated times after infection with 0.01 MOI of the indicated viruses. Data in **a** are geometric mean ± geometric SD. **b**, Sinonasal cavity from young BALB/c mice infected with 5000 PFU of SARS2-N501Y_MA30_, H&E stain (top panels). Regions of respiratory epithelium and olfactory epithelium exhibited uncommon regional scattered (bottom-left panel) to localized SARS-CoV-2 nucleocapsid immunostaining, (bottom panels). Scale bar 90 µm. OE: olfactory epithelium. **c**, Summary scores of nucleocapsid staining, as described in Methods (n=5). Data in **c** are mean ± s.e.m. **d**, Brains from uninfected or infected young BALB/c mice at 6 dpi lacked overt lesions, H&E stain. Scale bar 45 µm. **e**, Brains from uninfected mice or young BALB/c mice infected with 5000 PFU at 2,4, and 6 dpi revealed no SARS-CoV-2 nucleocapsid immunostaining. Scale bar 460 µm.

**Extended Data Figure 2. Histological analysis of extrapulmonary tissue**

Mice infected with 5000 PFU of SARS2-N501Y_MA30_ were sacrificed at 6 dpi with 5000 PFU of SARS2-N501Y_MA30_ and tissues were prepared for histological examination. Tissues from uninfected mice were analyzed in parallel. **a**–**e**, heart (**a**), liver (**b**), kidney (**c**), spleen (**d**), and small intestine (**e**) were studied. No overt, group-specific lesions were observed. Scale bars, 45 μm (**a, b, d**) and 90 μm (**c, e**), H&E stain. Two sections of each organ from 5 mice per group were evaluated.

**Extended Data Figure 3. Inflammatory mediators and immune effector cells in infected lungs**.

Mice were infected with 1000 PFU SARS2-N501Y_MA30_ to ensure survival until 6 dpi. **a**, Cytokine and chemokine transcripts were measured by qRT–PCR after isolation of RNA from the lungs of uninfected (0 dpi) and infected young BALB/c mice. Each lung was collected from an individual mouse. Mock (0 dpi), n=4; 2, 4, and 6 dpi, n=5. **b**, Quantification of immune cells (as gated in **e**) in the lungs (n = 3 for uninfected group; n = 4 and 5 for 4 and 6 dpi respectively) each lung was collected from an individual mouse). DC: dendritic cells; IMM: inflammatory monocytes/macrophages; PMN: polymorphonuclear cells. **c**, Representative FACS plots of IFNγ^+^/TNF CD4 and CD8 T cells (as gated in **e**) after stimulation with indicated peptide pools in the lungs of young BALB/c mice at 6 dpi. **d**, Summary data for IFNγ and TNF expression are shown (n=5). **e**, Gating strategy for identification of immune cells in lungs is shown. Data in **a, b, d** are mean ± s.e.m

**Extended Data Figure 4. Lymphopenia in SARS2-N501Y**_**MA30**_**-infected middle-aged mice**. Numbers of immune cells in the blood of young and middle-aged C57BL/6 mice at 3 and 6 days after infection with 5000 PFU of SARS2-N501Y_MA30_ (n=4 for uninfected controls and middle-aged mice at 3 dpi; n=5 for middle-aged mice at 6 dpi and young mice at 3 and 6 dpi). PMN: Polymorphonuclear cells. *P* values determined by two-tailed Student’s t test. *P* values indicated are determined from the corresponding time point compared to uninfected control of the same group. \**P*=0.0003 (3 dpi vs uninfected, young C57BL/6; CD45^+^ cells); \**P*=0.0194 (3 dpi vs uninfected, middle-aged C57BL/6, CD45^+^ cells); \**P*=0.0102 (6 dpi vs uninfected, middle-aged C57BL/6, CD45^+^ cells); \**P*=0.0046 (3 dpi vs uninfected, young C57BL/6; PMN); \**P*=0.0304 (3 dpi vs uninfected, middle-aged C57BL/6, PMN); \**P*=0.0318 (6 dpi vs uninfected, middle-aged C57BL/6, PMN); \**P*<0.0001 (3 dpi vs uninfected, young C57BL/6; B cells); \**P*=0.0264 (3 dpi vs uninfected, middle-aged C57BL/6, B cells); \**P*=0.0049 (6 dpi vs uninfected, middle-aged C57BL/6, B cells); \**P*=0.0025 (3 dpi vs uninfected, young C57BL/6; Total T cells); \**P*=0.0262 (3 dpi vs uninfected, middle-aged C57BL/6, Total T cells); \**P*=0.0051 (6 dpi vs uninfected, middle-aged C57BL/6, Total T cells); \**P*=0.0115 (3 dpi vs uninfected, young C57BL/6; CD4^+^ T cells); \**P*=0.0626 (3 dpi vs uninfected, middle-aged C57BL/6, CD4^+^ T cells); \**P*=0.0061 (6 dpi vs uninfected, middle-aged C57BL/6, CD4^+^ T cells); \**P*=0.0001 (3 dpi vs uninfected, young C57BL/6; CD8^+^ T cells); \**P*=0.0464 (3 dpi vs uninfected, middle-aged C57BL/6, CD8^+^ T cells); \**P*=0.0054 (6 dpi vs uninfected, middle-aged C57BL/6, CD8^+^ T cells). *\*P* ≤ 0.05, *\*\*P* ≤ 0.005, *\*\*\*P* ≤ 0.0005, *\*\*\*\*P* ≤ 0.0001. Data in **a** are mean ± s.e.m.

## REFERENCES

1 Channappanavar, R. & Perlman, S. Age-related susceptibility to coronavirus infections: role of impaired and dysregulated host immunity. J Clin Invest 130, 6204–6213, doi:10.1172/JCI144115 (2020).

2 Plante, J. A. et al. The variant gambit: COVID-19’s next move. Cell Host Microbe, doi:10.1016/j.chom.2021.02.020 (2021).

3 Vijay, R. et al. Critical role of phospholipase A2 group IID in age-related susceptibility to severe acute respiratory syndrome-CoV infection. J Exp Med 212, 1851–1868, doi:10.1084/jem.20150632 (2015).

4 Zhao, J., Zhao, J., Legge, K. & Perlman, S. Age-related increases in PGD(2) expression impair respiratory DC migration, resulting in diminished T cell responses upon respiratory virus infection in mice. J Clin Invest 121, 4921–4930, doi:10.1172/JCI59777 (2011).

5 Wan, Y., Shang, J., Graham, R., Baric, R. S. & Li, F. Receptor Recognition by the Novel Coronavirus from Wuhan: an Analysis Based on Decade-Long Structural Studies of SARS Coronavirus. J Virol 94, doi:10.1128/JVI.00127-20 (2020).

6 Bao, L. et al. The pathogenicity of SARS-CoV-2 in hACE2 transgenic mice. Nature 583, 830–833, doi:10.1038/s41586-020-2312-y (2020).

7 Jiang, R. D. et al. Pathogenesis of SARS-CoV-2 in Transgenic Mice Expressing Human Angiotensin-Converting Enzyme 2. Cell 182, 50–58 e58, doi:10.1016/j.cell.2020.05.027 (2020).

8 Winkler, E. S. et al. SARS-CoV-2 infection of human ACE2-transgenic mice causes severe lung inflammation and impaired function. Nat Immunol 21, 1327–1335, doi:10.1038/s41590-020-0778-2 (2020).

9 Zheng, J. et al. COVID-19 treatments and pathogenesis including anosmia in K18-hACE2 mice. Nature 589, 603–607, doi:10.1038/s41586-020-2943-z (2021).

10 Sun, S. H. et al. A Mouse Model of SARS-CoV-2 Infection and Pathogenesis. Cell Host Microbe 28, 124–133 e124, doi:10.1016/j.chom.2020.05.020 (2020).

11 Hassan, A. O. et al. A SARS-CoV-2 Infection Model in Mice Demonstrates Protection by Neutralizing Antibodies. Cell 182, 744–753 e744, doi:10.1016/j.cell.2020.06.011 (2020).

12 Sun, J. et al. Generation of a Broadly Useful Model for COVID-19 Pathogenesis, Vaccination, and Treatment. Cell 182, 734–743 e735, doi:10.1016/j.cell.2020.06.010 (2020).

13 Dinnon, K. H., 3rd et al. A mouse-adapted model of SARS-CoV-2 to test COVID-19 countermeasures. Nature 586, 560–566, doi:10.1038/s41586-020-2708-8 (2020).

14 Gu, H. et al. Adaptation of SARS-CoV-2 in BALB/c mice for testing vaccine efficacy. Science 369, 1603–1607, doi:10.1126/science.abc4730 (2020).

15 Leist, S. R. et al. A Mouse-Adapted SARS-CoV-2 Induces Acute Lung Injury and Mortality in Standard Laboratory Mice. Cell, doi:10.1016/j.cell.2020.09.050 (2020).

16 Peiris, J. S., Guan, Y. & Yuen, K. Y. Severe acute respiratory syndrome. Nat Med 10, S88–97 (2004).

17 Fehr, A. R. Bacterial Artificial Chromosome-Based Lambda Red Recombination with the I-SceI Homing Endonuclease for Genetic Alteration of MERS-CoV. Methods Mol Biol 2099, 53–68, doi:10.1007/978-1-0716-0211-9_5 (2020).

18 Starr, T. N. et al. Deep Mutational Scanning of SARS-CoV-2 Receptor Binding Domain Reveals Constraints on Folding and ACE2 Binding. Cell 182, 1295–1310 e1220, doi:10.1016/j.cell.2020.08.012 (2020).

19 Li, Q. et al. SARS-CoV-2 501Y.V2 variants lack higher infectivity but do have immune escape. Cell, doi:10.1016/j.cell.2021.02.042 (2021).

20 Wang, J. et al. Mouse-adapted SARS-CoV-2 replicates efficiently in the upper and lower respiratory tract of BALB/c and C57BL/6J mice. Protein Cell 11, 776–782, doi:10.1007/s13238-020-00767-x (2020).

21 Sun, H. et al. Characterization and structural basis of a lethal mouse-adapted SARS-CoV-2. bioRxiv, doi:doi: https://doi.org/10.1101/2020.11.10.377333 (2021).

22 Roberts, A. et al. A mouse-adapted SARS-coronavirus causes disease and mortality in BALB/c mice. PLoS Pathog 3, e5, doi:10.1371/journal.ppat.0030005 (2007).

23 Ackermann, M. et al. Pulmonary Vascular Endothelialitis, Thrombosis, and Angiogenesis in Covid-19. N Engl J Med 383, 120–128, doi:10.1056/NEJMoa2015432 (2020).

24 Galani, I. E. et al. Untuned antiviral immunity in COVID-19 revealed by temporal type I/III interferon patterns and flu comparison. Nat Immunol 22, 32–40, doi:10.1038/s41590-020-00840-x (2021).

25 Zhang, X. et al. Viral and host factors related to the clinical outcome of COVID-19. Nature 583, 437–440, doi:10.1038/s41586-020-2355-0 (2020).

26 Oslund, R. C., Cermak, N. & Gelb, M. H. Highly specific and broadly potent inhibitors of mammalian secreted phospholipases A2. J Med Chem 51, 4708–4714, doi:10.1021/jm800422v (2008).

27 Takahashi, G. et al. Effect of the potent and selective DP1 receptor antagonist, asapiprant (S-555739), in animal models of allergic rhinitis and allergic asthma. Eur J Pharmacol 765, 15–23, doi:10.1016/j.ejphar.2015.08.003 (2015).

28 Liu, Y. et al. The N501Y spike substitution enhances SARS-CoV-2 transmission. bioRxiv, doi:10.1101/2021.03.08.434499 (2021).

29 Marone, G. et al. Prostaglandin D2 receptor antagonists in allergic disorders: safety, efficacy, and future perspectives. Expert Opin Investig Drugs 28, 73–84, doi:10.1080/13543784.2019.1555237 (2019).

## Extended Data References

30 Yoshikawa, T. et al. Dynamic innate immune responses of human bronchial epithelial cells to severe acute respiratory syndrome-associated coronavirus infection. PLoS ONE 5, e8729, doi:10.1371/journal.pone.0008729 (2010).

31 Meyerholz, D. K. & Beck, A. P. Principles and approaches for reproducible scoring of tissue stains in research. Lab Invest 98, 844–855, doi:10.1038/s41374-018-0057-0 (2018).

32 Li, K. et al. Middle East Respiratory Syndrome Coronavirus Causes Multiple Organ Damage and Lethal Disease in Mice Transgenic for Human Dipeptidyl Peptidase 4. J Infect Dis 213, 712–722, doi:10.1093/infdis/jiv499 (2016).

33 Morin, A. et al. Collaboration gets the most out of software. Elife 2, e01456, doi:10.7554/eLife.01456 (2013).

