## Supplemental Figures for "Eicosanoid signaling as a therapeutic target in middle-aged mice with severe COVID-19"

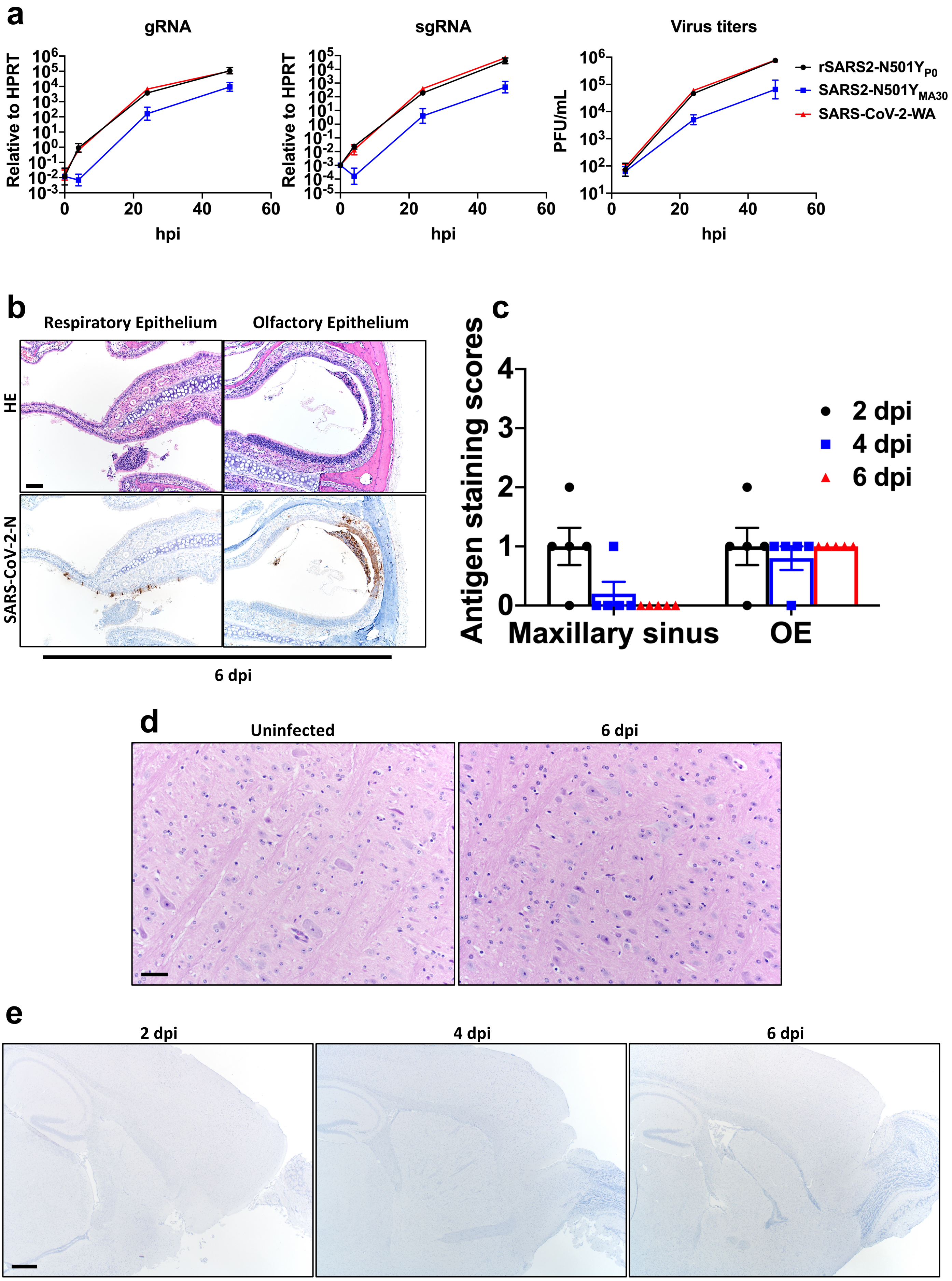


### Extended Data Figure 1

**Extended Data Figure 1. Infection of human respiratory cells and distribution of SARS2-N501Y$_{MA30}$ antigen in sinonasal cavity and brain.**

**a**, Quantification of genomic RNA (gRNA), subgenomic RNA (sgRNA) and virus titers in Calu-3 cells at the indicated times after infection with 0.01 MOI of the indicated viruses. Data in **a** are geometric mean ± geometric SD. **b**, Sinonasal cavity from young BALB/c mice infected with 5000 PFU of SARS2-N501Y$_{MA30}$, H&E stain (top panels). Regions of respiratory epithelium and olfactory epithelium exhibited uncommon regional scattered (bottom-left panel) to localized SARS-CoV-2 nucleocapsid immunostaining, (bottom panels). Scale bar 90 µm. OE: olfactory epithelium. **c**, Summary scores of nucleocapsid staining, as described in Methods (n=5). Data in **c** are mean ± s.e.m. **d**, Brains from uninfected or infected young BALB/c mice at 6 dpi lacked overt lesions, H&E stain. Scale bar 45 µm. **e**, Brains from uninfected mice or young BALB/c mice infected with 5000 PFU at 2,4, and 6 dpi revealed no SARS-CoV-2 nucleocapsid immunostaining. Scale bar 460 µm.

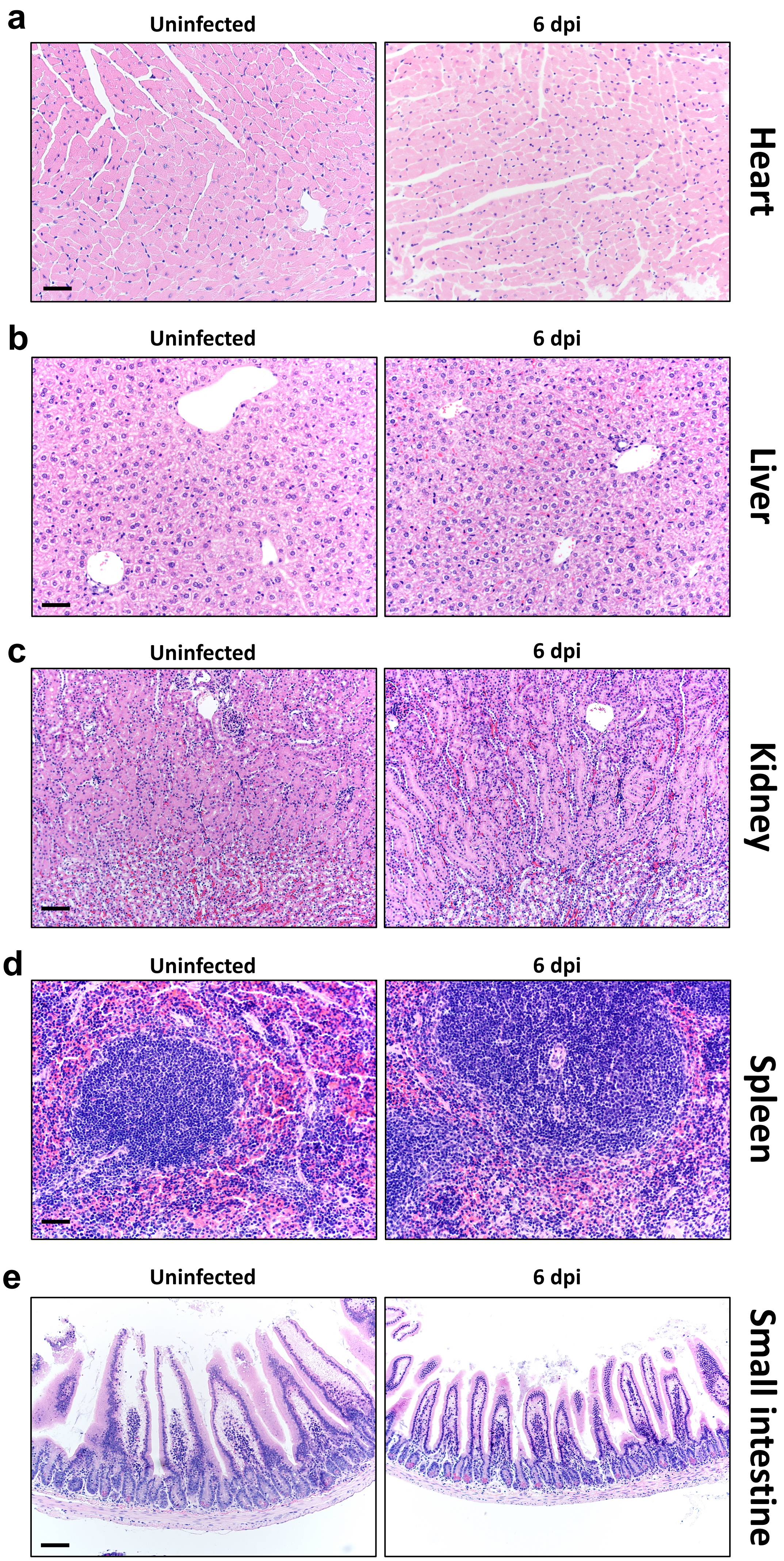


### Extended Data Figure 2

**Extended Data Figure 2. Histological analysis of extrapulmonary tissue**

Mice infected with 5000 PFU of SARS2-N501Y$_{MA30}$ were sacrificed at 6 dpi with 5000 PFU of SARS2-N501Y$_{MA30}$ and tissues were prepared for histological examination. Tissues from uninfected mice were analyzed in parallel. **a**–**e**, heart (**a**), liver (**b**), kidney (**c**), spleen (**d**), and small intestine (**e**) were studied. No overt, group-specific lesions were observed. Scale bars, 45 μm (**a, b**, **d**) and 90 μm (**c, e**), H&E stain. Two sections of each organ from 5 mice per group were evaluated.

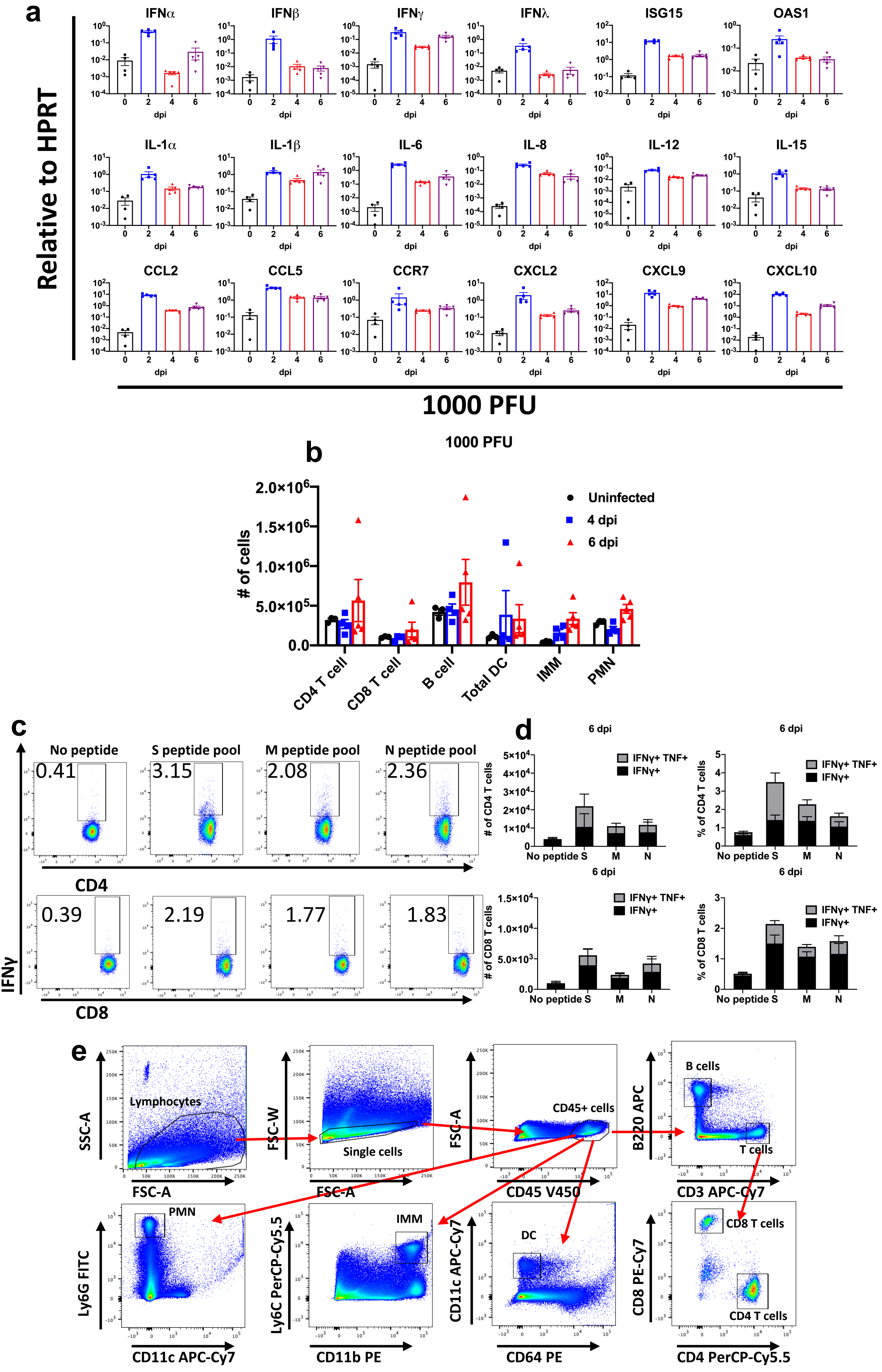


### Extended Data Figure 3

**Extended Data Figure 3. Inflammatory mediators and immune effector cells in infected lungs.**

Mice were infected with 1000 PFU SARS2-N501Y$_{MA30}$ to ensure survival until 6 dpi. **a,** Cytokine and chemokine transcripts were measured by qRT–PCR after isolation of RNA from the lungs of uninfected (0 dpi) and infected young BALB/c mice. Each lung was collected from an individual mouse. Mock (0 dpi), n=4; 2, 4, and 6 dpi, n=5. **b**, Quantification of immune cells (as gated in **e**) in the lungs (n = 3 for uninfected group; n = 4 and 5 for 4 and 6 dpi respectively) each lung was collected from an individual mouse). DC: dendritic cells; IMM: inflammatory monocytes/macrophages; PMN: polymorphonuclear cells. **c**, Representative FACS plots of IFNγ$^{+}$/TNF CD4 and CD8 T cells (as gated in **e**) after stimulation with indicated peptide pools in the lungs of young BALB/c mice at 6 dpi. **d**, Summary data for IFNγ and TNF expression are shown (n=5). **e**, Gating strategy for identification of immune cells in lungs is shown. Data in **a, b, d** are mean ± s.e.m

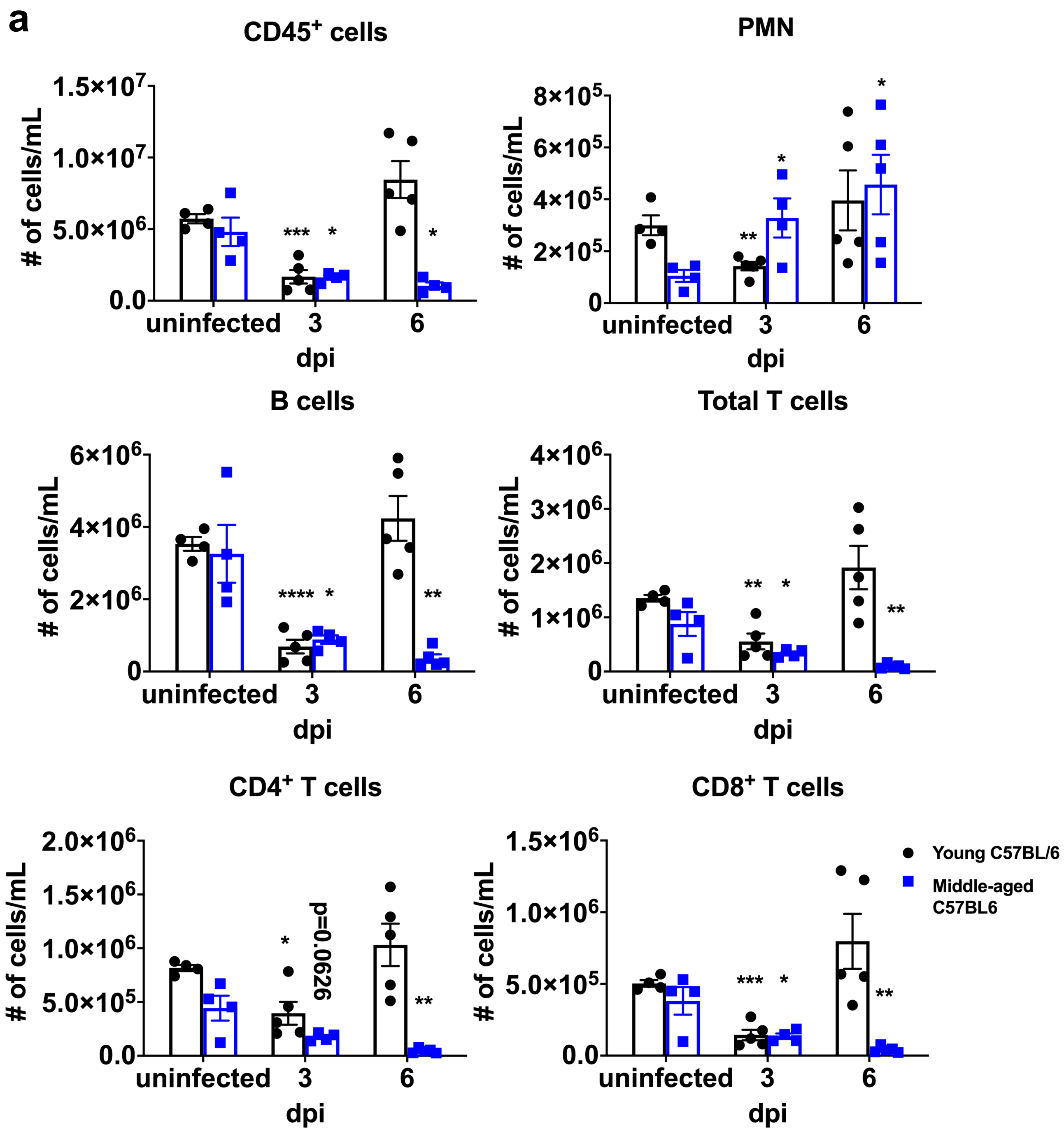


### Extended Data Figure 4

**Extended Data Figure 4. Lymphopenia in SARS2-N501Y$_{MA30}$-infected middle-aged mice.** Numbers of immune cells in the blood of young and middle-aged C57BL/6 mice at 3 and 6 days after infection with 5000 PFU of SARS2-N501Y$_{MA30}$ (n=4 for uninfected controls and middle-aged mice at 3 dpi; n=5 for middle-aged mice at 6 dpi and young mice at 3 and 6 dpi). PMN: Polymorphonuclear cells. *P* values determined by two-tailed Student's t test. *P* values indicated are determined from the corresponding time point compared to uninfected control of the same group. *$P$=0.0003 (3 dpi vs uninfected, young C57BL/6; CD45$^+$ cells); *$P$=0.0194 (3 dpi vs uninfected, middle-aged C57BL/6, CD45$^+$ cells); *$P$=0.0102 (6 dpi vs uninfected, middle-aged C57BL/6, CD45$^+$ cells); *$P$=0.0046 (3 dpi vs uninfected, young C57BL/6; PMN); *$P$=0.0304 (3 dpi vs uninfected, middle-aged C57BL/6, PMN); *$P$=0.0318 (6 dpi vs uninfected, middle-aged C57BL/6, PMN); *$P$<0.0001 (3 dpi vs uninfected, young C57BL/6; B cells); *$P$=0.0264 (3 dpi vs uninfected, middle-aged C57BL/6, B cells); *$P$=0.0049 (6 dpi vs uninfected, middle-aged C57BL/6, B cells); *$P$=0.0025 (3 dpi vs uninfected, young C57BL/6; Total T cells); *$P$=0.0262 (3 dpi vs uninfected, middle-aged C57BL/6, Total T cells); *$P$=0.0051 (6 dpi vs uninfected, middle-aged C57BL/6, Total T cells); *$P$=0.0115 (3 dpi vs uninfected, young C57BL/6; CD4$^+$ T cells); *$P$=0.0626 (3 dpi vs uninfected, middle-aged C57BL/6, CD4$^+$ T cells); *$P$=0.0061 (6 dpi vs uninfected, middle-aged C57BL/6, CD4$^+$ T cells); *$P$=0.0001 (3 dpi vs uninfected, young C57BL/6; CD8$^+$ T cells); *$P$=0.0464 (3 dpi vs uninfected, middle-aged C57BL/6, CD8$^+$ T cells); *$P$=0.0054 (6 dpi vs uninfected, middle-aged C57BL/6, CD8$^+$ T cells). *$P \leq 0.05$, **$P \leq 0.005$, ***$P \leq 0.0005$, ****$P \leq 0.0001$. Data in **a** are mean ± s.e.m.
